# Genome analysis of *Salmonella enterica* serovar Typhimurium bacteriophage L, indicator for StySA (StyLT2III) restriction-modification system action

**DOI:** 10.1101/2020.10.05.325894

**Authors:** Julie Zaworski, Colleen McClung, Cristian Ruse, Peter R. Weigele, Roger W. Hendrix, Ching-Chung Ko, Robert Edgar, Graham F. Hatfull, Sherwood R. Casjens, Elisabeth A. Raleigh

**Author notes:** Deceased (Roger W. Hendrix). Corresponding author: Elisabeth A Raleigh, Research Department, New England Biolabs, 240 County Road, Ipswich, MA 01938.

## Abstract

Bacteriophage L, a P22-like phage of *Salmonella enterica* sv Typhimurium LT2, was important for definition of mosaic organization of the lambdoid phage family and for characterization of restriction-modification systems of *Salmonella*. We report the complete genome sequences of bacteriophage L *cI*^−^40 *13*^−^*am*43 and L *cII*^−^101; the deduced sequence of wildtype L is 40,633 bp long with a 47.5% GC content. We compare this sequence with those of P22 and ST64T, and predict 71 Coding Sequences, 2 tRNA genes and 14 intergenic rho-independent transcription terminators. The overall genome organization of L agrees with earlier genetic and physical evidence; for example, no secondary immunity region (*ImmI*: *ant*, *arc*) or genes for superinfection exclusion (*sieA* and *sieB*) are present. Proteomic analysis confirmed identification of virion proteins, along with low levels of assembly intermediates and host cell envelope proteins. The genome of L is 99.9% identical at the nucleotide level to that reported for phage ST64T, despite isolation on different continents ~35 years apart. DNA modification by the epigenetic regulator Dam is generally incomplete. Dam modification is also selectively missing in one location, corresponding to the P22 phase-variation-sensitive promoter region of the serotype-converting *gtrABC* operon. The number of sites for SenLTIII (StySA) action may account for stronger restriction of L (13 sites) than of P22 (3 sites).

## INTRODUCTION

Bacteriophage L was isolated in 1967 from *S. enterica* sv Typhimurium LT2(L) after UV light or mitomycin induction (Bezdek and Amati 1967, Bezdek and Amati 1968). We note that the LT2 reference strain ATCC700720 (McClelland, Sanderson et al. 2001) and common laboratory derivatives do not carry this prophage. However, in early classification schemes dependent on phage sensitivity patterns, numerous independent natural *Salmonella* Typhimurium isolates would be defined as “LT2” (e.g. (Zinder and Lederberg 1952)). Whatever its history, phage L virion structure, properties and gene organization are similar but not identical to those of P22 (Bezdek and Amati 1967, Bezdek and Amati 1968, Bezdek, Soska et al. 1970, Susskind, Botstein et al. 1974, Susskind, Wright et al. 1974, Susskind and Botstein 1978, Hilliker 1981), as is its sequence arrangement based on DNA hybridization, restriction analysis (Hilliker 1981, Hayden, Adams et al. 1985) and partial sequence studies (Sauer, Krovatin et al. 1983, Schicklmaier and Schmieger 1997, Gilcrease, Winn-Stapley et al. 2005). Like P22, it is a short-tailed, temperate, dsDNA bacteriophage that forms turbid plaques. The similarity of virion structure and gene organization was important in the elucidation of the second immunity region of P22 (*immI, mnt-arc-ant*) (Bezdek and Amati 1968, Susskind and Botstein 1978, Sauer, Krovatin et al. 1983, Schicklmaier and Schmieger 1997), and in characterization of the superinfection exclusion mechanisms (*sieA* and *sieB*) of P22 (Susskind, Botstein et al. 1974, Susskind, Wright et al. 1974). The close relationship between L and P22 was known genetically through the ability of the two phages to form viable hybrids (Bezdek and Amati 1967, Bezdek and Amati 1968, Favre, Amati et al. 1968, Susskind and Botstein 1978) and by complementation of P22 mutants by mixed infection (Kahmann and Prell 1971) or by an L prophage (Schicklmaier and Schmieger 1997). Physically, DNA:DNA hybridization and/or complementation has been demonstrated between L and P22 regulatory genes *c3, 23*, and *24*, virion assembly genes *1, 2, 3, 5, 8, 9, 10, 16*, and *20*, and lysis genes *13, 15*, and *19* (Wiggins and Hilliker 1985, Schicklmaier and Schmieger 1997). On the other hand, its CII (ortholog of P22 C2 and λ CI) and Cro repressors and CI (ortholog of P22 C1 and λ CII) establishment activator proteins have different specificities from those of P22 (2, 10). The mosaicism of the late operons of the P22-like bacteriophages was reviewed in 2011 (Casjens and Thuman-Commike 2011).

Phage L was also used together with P22 and P3 to study restriction-modification (RM) systems in *S. enterica* sv Typhimurium LT2 and LT7 (Colson and Colson 1971, Colson and Van Pel 1974, Hattman, Schlagman et al. 1976, Bullas and Ryu 1983, Fuller-Pace, Bullas et al. 1984). These prokaryotic defense system components are characterized by two activities: protective modification of the host DNA at specific recognition sites, and restriction action, which interferes with establishment of unmodified DNA entering the cell. This permits the host to distinguish self from nonself DNA. Phage L was a specific tester for the StySA restriction-modification system in *S. enterica* sv Typhimurium LT2 and LT7 (Colson and Colson 1971, Colson and Van Pel 1974, Hattman, Schlagman et al. 1976, Bullas and Ryu 1983).

The complete circularized L genome sequence and annotations presented here confirm reported similarities between P22 (Vander Byl and Kropinski 2000, Pedulla, Ford et al. 2003) and L (Sauer, Krovatin et al. 1983, Schicklmaier and Schmieger 1997, Gilcrease, Winn-Stapley et al. 2005) and is compared with ST64T (Mmolawa, Schmieger et al. 2003). Virion proteins are also characterized.

## MATERIALS AND METHODS

### Phages - growth, purification, storage and DNA isolation

Phages L *cl*^−^40, *13*^−^*am*43 (Bode 1979)and L *cII*^−^101 (Susskind and Botstein 1978) were the kind gifts of W. Bode and D. Botstein, respectively. They were propagated from single plaques in *Salmonella* strain DB7000 (Winston, Botstein et al. 1979) or STK005 (an isolate of L5000 (Bullas and Ryu 1983)). Virions were purified by cesium chloride step gradient centrifugation as described (Earnshaw, Casjens et al. 1976). Phage lysates were prepared by addition of 0.01% v/v of chloroform to the culture 90-120 min after infection, cell debris were removed by centrifugation and the supernatant was stored at 8°C.

Genomic DNA was extracted from virions using a phenol-chloroform extraction protocol. Briefly, 200 μl of phage particles were mixed with 200 μl TE (25 μl 10% SDS, 50 μl 1 M Tris-HCl pH 8.0, 25 μl 0.5M EDTA pH 8.0) and 200 μg protease K, incubated for 20 min at 56-65°C, and 500 μl phenol:chloroform:isoamyl alcohol (25:24:1) was added. The tube was shaken for 2 minutes and centrifuged. The top aqueous layer was mixed with one volume of 100% chloroform, shaken and centrifuged. This step was repeated once. gDNA was concentrated using ethanol precipitation and resuspended in TE buffer (Casjens and Gilcrease 2009).

### Sequence determination

The phage L *c*I^−^40 *13*^−^*am*43 genome was sequenced at New England Biolabs by combining data from Illumina and PacBio RS2 methods. An Illumina DNA library was prepared with the NEBNext^®^ Ultra™ II FS DNA Library Prep Kit (NEB, Ipswich, USA) and multiplex barcoded with NEBNext^®^ Multiplex Oligos for Illumina^®^ (Index Primers Set 2) (NEB, Ipswich, USA). An Illumina MiSeq device (Illumina Inc., San Diego, CA, USA) was used to generate 75-bp pair-end reads. These reads were trimmed using Trim Galore version 0.5.0 with adapter sequence ‘AGATCGGAAGAGC’. Short reads under 25 bp and unpaired reads were discarded. For long read sequencing, a SMRTbell library was constructed from a genomic DNA sample sheared to an average size of ~10 kb using the G-tubes protocol (Covaris, Woburn, MA, USA). Single strands were removed, ends repaired, and fragments ligated to hairpin adapters. Incompletely formed SMRTbell templates were eliminated by digestion with a combination of Exonuclease III and Exonuclease VII (New England BioLabs; Ipswich, MA, USA). Sequencing employed the PacBio RS2 instrument using the DNA/Polymerase Binding Kit P4, MagBead Loading Kit, and Sequencing Kit 2.0 (all Pacific Biosciences). Data from 1 SMRT cell was used, with 240 mins movie per cell. *De novo* PacBio assembly was performed with Canu 1.7.1 (Koren, Walenz et al. 2017) on 4,081 generated Circular Consensus Sequence (CCS) with RS_ReadsOfInsert (Minimum full passes = 5 and Minimum predicted accuracy = 95) from the PacBio SMRT portal. One contig of length 46,371 bp was obtained. This contig was circularized using Circlator (Hunt, Silva et al. 2015) to obtain a 40,663 bp closed genome. PacBioRS2 CCS reads were combined with the trimmed Illumina reads, to perform a hybrid *de novo* assembly using Unicycler (Wick, Judd et al. 2017). One 40,663 bp complete circular contig was thus obtained with a coverage depth of 2,690.67x that aligned with the PacBioRS2 *de novo* assembly with 100% identity. PacBio reads were used to detect m6A-modified DNA motifs with SMRT motif and modification analysis version 2.3.0 as described in (Clark, Murray et al. 2012).

The phage L *cII*^−^101 genome was sequenced at the University of Pittsburgh by dideoxy chain termination methods as described (Pedulla, Ford et al. 2003) and at New England Biolabs by Illumina techology (above). In Pittsburgh, sequencing of a plasmid DNA library was performed to an average depth of 7-fold coverage. The data assembled into a single circular contig with PhredPhrap (Gordon, Abajian et al. 1998). At New England Biolabs, the Illumina sequencing and trimming were performed as for phage L *c*I^−^40 *13*^−^*am*43. The 2,313,187 raw pair-end trimmed reads remaining (97% of the reads) were mapped to the L *c*I^−^40 *13*^−^*am*43 reference genome generated (above) with Bowtie2 (Langmead and Salzberg 2012), sorted and indexed with SAMtools (Li, Handsaker et al. 2009). The two methods gave identical 40,664 bp circular sequences. The phage L virion chromosome is known to be terminally redundant and circularly permuted so a circular sequence assembly is expected (Hayden, Adams et al. 1985).

### Gene identification and nomenclature

Annotation of the L genome features was performed with the Rapid Annotations using Subsystems Technology (Aziz, Bartels et al. 2008) and PHASTER, an upgraded version of PHAge Search Tool (PHAST) (Zhou, Liang et al. 2011, Arndt, Grant et al. 2016) servers, which predicted respectively 66/1/0 and 64/0/2 coding sequences (CDSs)/tRNAs/attachment sites, followed by manual curation. Among these annotations, we retained the 58 features predicted by both programs with the same coordinates, 7 predicted by either PHAST or RAST but corresponding to other phage annotations. Five annotations were added manually (*gtrA, xis, ninA, ninD* and *rz1*) from comparisons with phages ST64T (AY052766), P22 (TPA:BK000583) and other phages (Supplementary data). The gene coding the protein Dec annotation was added based on a previous report (Gilcrease, Winn-Stapley et al. 2005). The tRNA genes were predicted using tRNAscan-SE online tool (Lowe and Chan 2016, Chan and Lowe 2019) (Table S1).

Because of the long history of study of P22 and related phages, a considerable body of functional assignment has already accumulated. To maintain consistency with that literature, in this work we used names derived from P22 for L ORFs where sequence homology exists. Other genes were given the same name as their homologue in phage ST64T (see Results) or other phages. For *immC* genes, we have used the Roman numeral nomenclature of Bezdek *et al*. (Bezdek and Amati 1967, Bezdek and Amati 1968, Bezdek, Soska et al. 1970, Susskind, Botstein et al. 1974, Susskind, Wright et al. 1974, Susskind and Botstein 1978, Hilliker 1981), rather than P22 Arabic numbers: repressor of lysogeny is *cII* (RAST assigns “repressor”); activator of *cII* expression is *cI* (RAST assigns “transcriptional activator”).

### Proteomic analysis

The purified virion preparation (buffer: 10 mM TrisCl PH7.5, 1 mM MgCl2) was digested with trypsin using a FASP protocol (Filter Aided Sample Prep from Expedeon) (Wisniewski, Zougman et al. 2009). The resulting peptides were chromatographically separated over a reversed phase C18 column (via Proxeon Easy nLC II), eluted with a gradient of 10-45% acetonitrile and subjected to mass spectrometer (Thermo LTQ Orbitrap XL) LC-MS/MS analysis. The sample was run in triplicate.

Data were analyzed with PEAKS and Proteome Discoverer 2.4 (Bioinformatics Solutions, Inc.) to identify virion proteins. Peptide spectrum matches (spectral counting) were further processed to calculate cNSAF (Mosley, Sardiu et al. 2011). Abundances of protein signals were quantified using TopN peptides analysis of peak areas. Peptide and protein grouping validation was set at 1% FDR. Composition of virion was thus estimated with two label free quantitative approaches: spectral counting (MS2) and TopN method (MS1) (Krey, Wilmarth et al. 2014). Table S2 in the supplementary data contains the detailed results of the analysis.

### Data availability

Strains and variants are available upon request. Raw Pacific Bioscience RSII reads (SRR12424739) and Miseq Illumina raw reads (SRR12424740 and SRR12424741) have been deposited in the NCBI Bioproject PRJNA605961, and the raw dideoxy chain termination sequencing data for L *cII*^−^101 has been deposited in the NCBI SequenceReadArchive (SRA) with accession number SRR8384267. The phage L genome sequence and annotations have been deposited in GenBank (accession L_cI-40_13-am43 MW013502 and L_cII-101 MW013503). Table S1 and S2 supplementary files are available on GSA Figshare portal. Table S1 in supplementary data contains the refined annotations, including name, type, inference method, start, stop, length, direction, sequence, putative product and translation. Table S2 contains the proteomic analysis spectral counts and protein IDs.

## RESULTS AND DISCUSSION

### The phage L genome sequence

Two phage L genomes, L *cII*^−^101 and L *cI*^−^40 *13*^−^*am*43, were completely sequenced as described in MATERIALS AND METHODS and found to be 40,664 bp and 40,663 bp long, respectively. Note that the difference in length of the two genomes (above) is due to the *cII*^−^*101* 1 bp insertion. Previously determined phage L sequences match 10,526 bp: 8,338 bp of the head gene region ((Gilcrease, Winn-Stapley et al. 2005), AY795968) and 2,188 bp of the *immunity C* region ((Schicklmaier and Schmieger 1997); X94331). The head gene region sequence is identical to the corresponding region of our complete genome sequences. However, the previous *immunity C* sequence contains a 25 bp deletion and 11 other bps that differ from our sequences. These differences are likely sequencing errors, since our three independent determinations are identical at these locations. Thus, our sequence corrects errors in the previously reported sequences of L genes *24, cro, cI* and *cII* (below). There are only 8 single bp differences between our L *cII* 101 and L *cI* 40 *13 am*43 genome sequences (listed in Table 1). Comparison of these two L genomes, the previously reported sequence of the L *cII* gene (Schicklmaier and Schmieger 1997), and homologous genes in P22 allowed three of these differences to be unambiguously identified as single bp changes due to the three known mutations in these L strains: *cII*^−^ 101 is a 1 bp insertion frameshift in the prophage repressor gene; *cI*^−^40 is a missense mutation in this transcriptional activator gene; and *13*^−^*am*43 creates a nonsense codon in the holin gene (Table 1).

**Table 1:** Nucleotide sequence differences among phage L strains and phage ST64T. Column 1: coordinate of the mutations in *L cI 40 13am43*. Column 2, 3 and 4: nucleotide at the coordinate in L cI 40 13am43, L cII 101 and ST64T genomes respectively. Column 4: gene annotation in which the mutation is located. *Column 5:* consequences of the observed mutations.

| **L cl 40**<br>**13am43**<br>**coordinate** | **L cl 40**<br>**13am43**<br>**nucleotide** | **L cII 101**<br>**nucleotide** | **ST64T**<br>**nucleotide** | **L Gene**<br>**(function)** | **Difference details** |
| --- | --- | --- | --- | --- | --- |
| 6512 | A | G | G | orf186 | Difference in codon 38; Ile in L cl 40 13am43, Val in L cII 101 |
| 21362 | A | G | G | eaC | synonymous difference in codon 193 |
| 23697 | A | G | G | orf87 | synonymous difference in codon 84 |
| 24146 | A | G | G | orf91 | synonymous difference in codon 23 |
| 24823 | ΔT | ΔT | T | orf109 | one T insertion frameshift in codon 59 |
| 27443 | A | G | G | 17 | synonymous difference in codon 96 |
| 28342 | Δ | Δ | 15 bp | orf232 | ST64T has one more 15 bp repeat than L (15 imperfect 5 AA repeats in ST64T) |
| 30260 | – | C insert | – | cII<br>(repressor;<br>RAST) | 1 C insertion in run of C's at codon 46 is the *cII* 101 frameshift mutation |
| 30877 | G | A | A | cl<br>(activator) | G is *cl* 40 mutation in codon 27 (Gln to Arg) |
| 38322 | T | C | C | 13 (holin) | T is *amber* mutation in codon 50 Gln (CAG to TAG) |

The five other single bp differences between these two phage genomes are synonymous single bp differences in *eaC, orf87, orf91* and gene *17*, and an isoleucine/valine coding difference in *orf186*. The fact that for each of these five differences the L *cII*^−^101 has the same bp as phage ST64T (Table 1 and see below) is consistent with the idea that they were generated in L *cI*^−^40 *13*^−^*am*43 by mutagenesis that was very likely used to isolate the *amber* mutation and that the L *cII*^−^101 sequence represents the L wild type sequence at these locations. The resulting genome deduced for the inferred wild type L is 40,633 bp long with a 47.5% G+C content. This length agrees very well with the estimate previously obtained by restriction fragment analysis, 40,650 ± 400 bp (Hayden, Adams et al. 1985).

To further assess the accuracy of the circularized assembly, restriction digests of the phage genomic DNA by SapI, PvuII and EcoRI were performed, and the resulting fragment length measurements agree perfectly the chromosome structure obtained *in silico*, Figure 1. L *cI*^−^40 *13*^−^*am*43 virions each contain a terminal redundancy of ~6.5% (~2.5 kb), since more than a unit genome is packaged from long concatemers beginning at a site within gene *3* (a headful packaging mechanism (Hayden, Adams et al. 1985)), and because of this headful packaging, the fragment containing the 3’ portion of gene *3* is submolar and variable (starred fragments in Figure 1) (Casjens and Gilcrease 2009).

**Figure 1.**
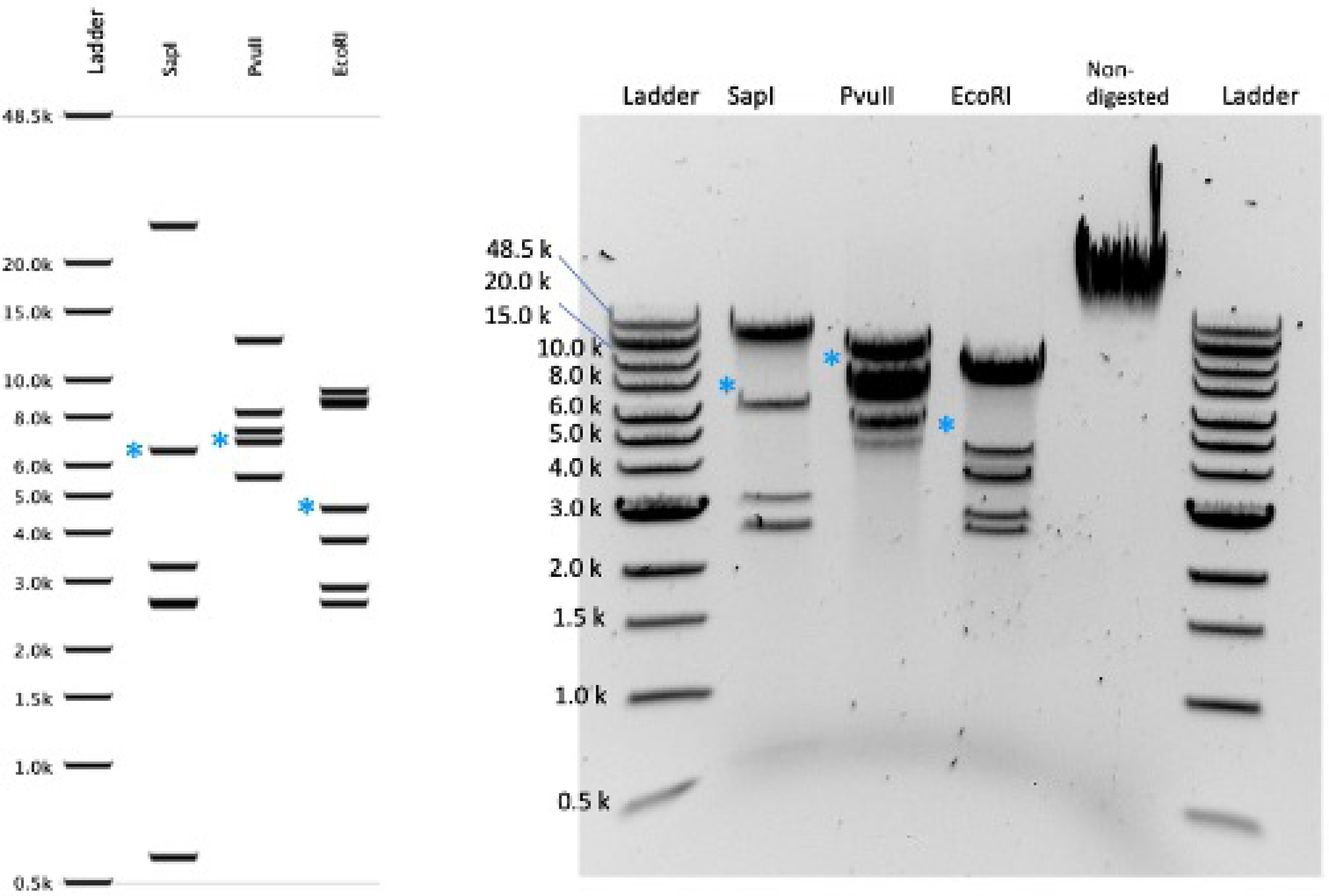
Restriction analysis of the L genome. Right panel: L genome structure was verified by restriction digestion analysis as follows: 1 ug of DNA in a 50 ul reaction was incubated for 1 h at 37°C with each of three enzymes separately (EcoRI-HF, SapI and PvuII-HF; New England Biolabs), and run on a 0.8% agarose electrophoresis gel with the 1 kb Extend Ladder (New England Biolabs). Asterisks (*) denote fragments with one end cut by the headful packaging. Left panel: *in silico* fragment predictions for the circular genome).

We identified 71 protein-encoding genes in the L genome, 48 of which have homologs in the well-studied phage P22. In addition, two tRNA genes are present. The L genes are discussed below in more detail.

### Similarities among the genomes of phages L, ST64T and P22

Previous studies have shown that phage L is a member of the P22-like subgroup of lambdoid phages. In particular, comparison of P22 and L by restriction mapping (10, 24,34) and heteroduplex analysis (Wiggins and Hilliker 1985) have indicated that they have substantial regions of nucleotide sequence similarity and our sequence shows high similarity in those regions. Thus, comparison with the well-studied phage P22 (sequenced and corrected in (Vander Byl and Kropinski 2000, Pedulla, Ford et al. 2003)) is informative in understanding the L genome, and this is done in the following section.

We also note that the inferred wildtype L genome (above) and the phage ST64T genome (AY052766) (Mmolawa, Schmieger et al. 2003) are extremely similar, with >99.9% nucleotide sequence identity. Of the 71 annotated L genes, 61 show 100% sequence identity with ST64T homologs. A total of 16 bp differences occur at only two sites (listed in Table 1), a one bp deletion in L affects orf109 and a 15 bp deletion removes one of the pentapeptide repeats in the L early left operon gene *orf232* without affecting the reading frame. This extremely high similarity between the independently isolated L and ST64T phages is very unusual given the tremendous diversity of the P22-like phages, but in spite of the murky history of phage L, it seems unlikely that the two phages share any laboratory history.

### Phage L predicted genes – conserved and variable segments and specificity divergence

In the following sections, the phage L genes are discussed from left to right across the genome map shown in Figure 2.

**Figure 2.**
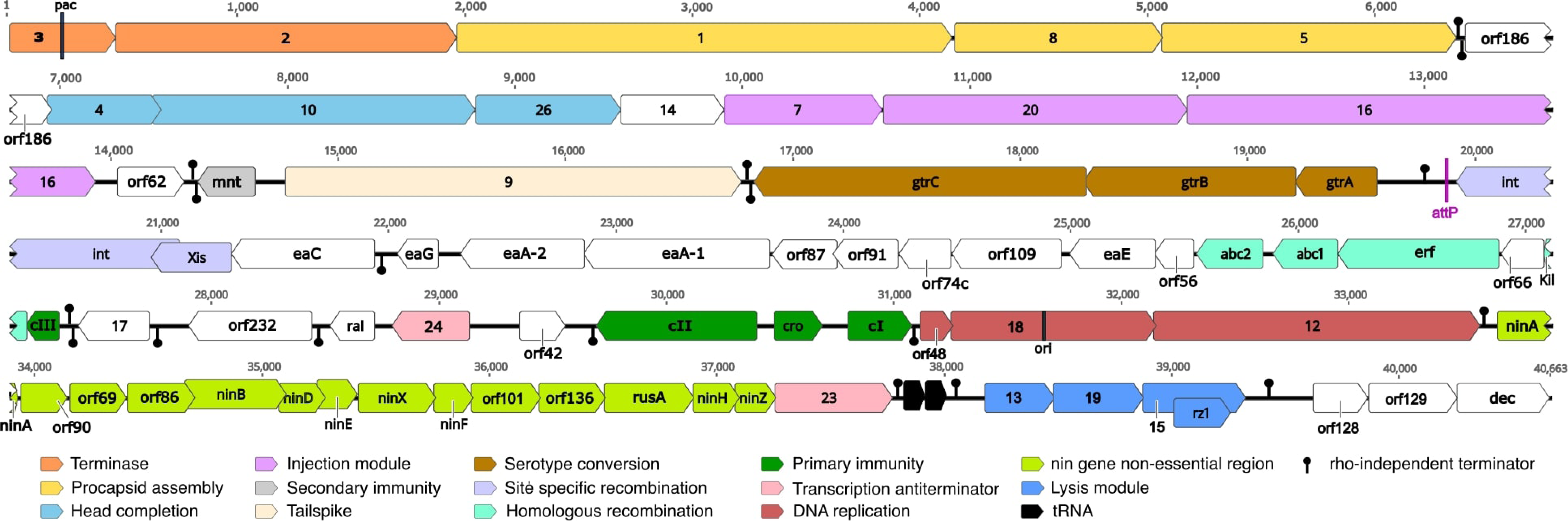
Phage L genome organization. Linear representation of the genome. The CDS are represented by arrows, with color differentiating functional modules or regions. tRNA positions are marked by black arrows. Transcription terminators were predicted *in silico*; those blocking rightward are above the line and those blocking leftward transcription are below the line.

#### Virion assembly genes

By analogy with phages λ and P22, the phage L virion assembly genes are almost certainly expressed from a single late promoter and so are part of the late operon (see below). Overall, the L virion assembly genes are similar to those of P22 except for an extra major phage L virion decoration protein, the product of the *dec* gene, 60 trimers of which are bound to the exterior surface and stabilize the virion (Gilcrease, Winn-Stapley et al. 2005, Tang, Gilcrease et al. 2006). The first large homologous region runs from gene 3 to gene 14 (Figure 3). The DNA packaging/procapsid assembly gene module is highly conserved with 95.1 % identity from position 0 to 6,360. This segment contains the terminase genes (*3* and *2* in orange, Figures 2 and 3) and procapsid assembly genes (*1, 8* and *5* in yellow, Figure 2 and 3). These were all shown to be functionally interchangeable with those of P22 in complementation experiments (12). The 22 bp sequence that P22-like phages use to initiate DNA packaging, called *pac*, lies inside gene *3* (Leavitt, Gilcrease et al. 2013); L gene *3* contains all the critical bp of the P22 *pac* site at bp 268-264 (Wu, Sampson et al. 2002), Figure 2. Thus, L very likely has the same DNA packaging specificity as P22.

**Figure 3.**
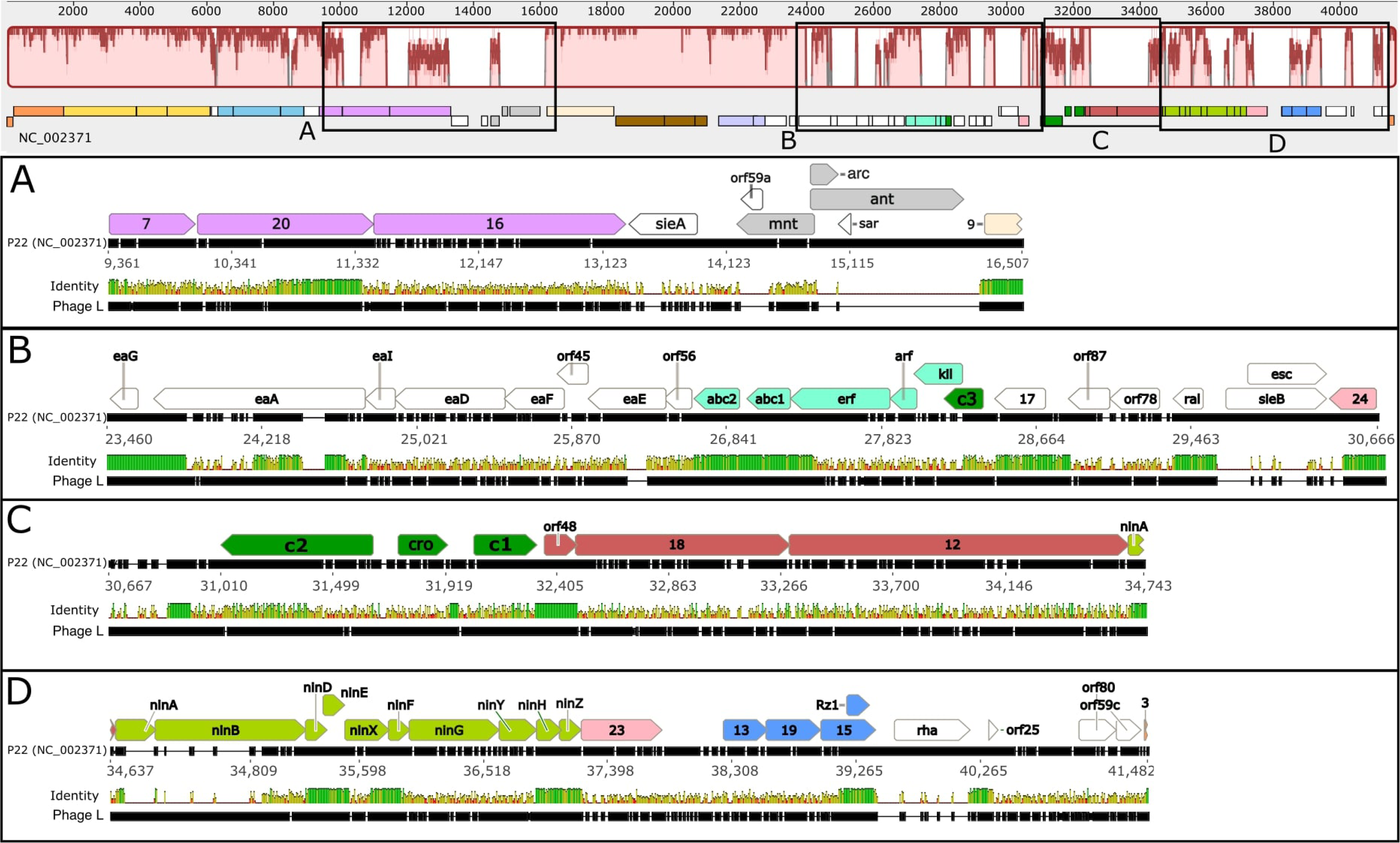
Comparison of phage L and P22 genomes. The top panel displays the Mauve alignment of whole L and P22 genomes with coding sequences of P22 represented below. Pink regions of the Mauve alignment correspond to significantly similar stretches while white regions correspond to low homology areas. Insets A, B, C and D display MAFFT-aligned closeups of low homology areas boxed above. For each inset, top row: the annotated CDSs of P22 are aligned with the second row, P22 reference sequence; third row, nucleotide identity between P22 and L; fourth row: L reference sequence (determined here). Black ticks in each genome reference row correspond to aligned segments, white ticks to gaps. The identity row is vertically scaled to the degree of nucleotide identity of the aligned sequences. 100% identity is dark green, chartreuse is shorter with lower identity, and red represents the lowest identity, including gapped regions.

The head completion module (light blue) comprising genes *4, 10* and *26* extends the high synteny region interrupted by *orf186* (detailed function unknown; white), with protein identities 97, 95.8 and 94%, respectively. Genes *7, 20* and *16* (protein identities 65.2%, 62% and 29.8%) comprise the injection protein module (pink), with patchier and overall lower DNA similarity to P22 (Figure 3A and Supplementary data). Such patchy similarity is frequent with this group of phages (Casjens and Thuman-Commike 2011); here we note that L and P22 have a section of high similarity at the 3’ end of gene *20*.

The tailspike proteins (gene *9*) of P22 and L are 98.7% identical, so it is not surprising that they have been shown to cross-react immunologically (Bezdek and Amati 1967) and are functionally interchangeable (Schicklmaier and Schmieger 1997). Their near identity suggests that they almost certainly adsorb to the same Typhimurium O-antigen polysaccharide, and this is consistent with L’s ability to infect Typhimurium strains such as LT2.

#### Secondary Immunity

L-P22 hybrids were used to define the P22 *immI* region (light grey in Figure 2 and Figure 3A) (Susskind and Botstein 1978). This region between genes *16* and *9*, inside the late operon but largely not expressed from the late operon message, is very variable in the P22-like phages (see Figure 3 of (Villafane, Zayas et al. 2008)). In P22 it contains the *immI* region that includes genes for an antirepressor gene *ant* and genes *mnt* and *arc* that control its expression, as well as the superinfection exclusion gene *sieA* (which is thought to prevent injection of DNA by other related phages (Susskind, Botstein et al. 1974)). This 836 bp phage L region lacks *sieA, ant* and *arc* but carries a rather poorly conserved homolog of *mnt* and an apparent short gene, *orf62*, of unknown function (Figure 3A). The *mnt*-like gene could have acquired another function in L or be a remnant in a degrading *immI* region. L’s lack of the above genes is consistent with genetic evidence that phage L lacks a functional *immI* region (Susskind and Botstein 1978) and with previous DNA hybridization studies (Hayden, Adams et al. 1985, Schicklmaier and Schmieger 1997)

#### Serotype conversion module

The phage P22 prophage is able to modify the host O-antigen through the action of its *gtrABC* genes; these are highly homologous to *gtr* genes of phages SfV, SfII, and SfX (Adhikari, Allison et al. 1999, Broadbent, Davies et al. 2010). The L genome sequence contains *gtrABC* genes that are nearly identical to those of P22 - *gtrA*, 100%; *gtrB*, 99.7% and *gtrC*, 98.8% (brown in Figure 2 and 3) - so an L prophage would be expected to convert the O12 antigen of its *S*. Typhimurium host to the O1 serotype (Broadbent et al, 2010). In P22, the *gtr* operon is subject to phase variation that is mediated by binding of the regulator host protein OxyR and action of Dam methylase (Broadbent, Davies et al. 2010). ST64T was shown to carry out serotype conversion by immunologic methods (Mmolawa, Schmieger et al. 2003), but its regulation was not investigated. Several sequence differences between L and P22 are present just upstream of the *gtrABC* operon, but the Dam methylation sites, OxyR binding sites and their spacing appear to be the same in both phages, so phase-variable expression in the L prophage is likely to be the same as that in P22. Note that these regulatory Dam sites are undermethylated in the virion (see below, DNA Methylation and Restriction).

#### Site-specific recombination (integration) module

The site-specific recombination module *int* and *xis* genes (lavender in figure 2) are also very similar to those of P22 with 98.4% and 100% protein identity, respectively. Although the phage L integration site in the host chromosome has not been experimentally determined, the P22 *attP* site “ATGCGAAGGTCGTAGGTTCGACT” (Lindsey, Martínez et al. 1992) is present at bp 19,865-19,887 at the expected location in the L sequence, inside a false tRNA gene (annotated at L bp 19,823-19,903) that lies just downstream of the *int* gene and rebuilds an intact *thrW* gene at one end of the prophage upon integration (Lindsey, Martínez et al. 1992). These very high similarities with P22 make it essentially certain that L integrates at the P22 *attB* site in the *thrW* tRNA gene.

#### The early left operon

The L early left operon contains 20 genes, a number of which are similar to those of P22 (Figure 3B). Known functions for homologs in other phages are as follows: homologous recombination (*abc1, abc2* and *erf*), host killing by blocking cell division (*kil*), relief of type I restriction (*ral*) and establishment of lysogeny (*cIII*). However, for a majority of these genes, functions are not known. L lacks the superinfection exclusion gene *sieB* carried here in P22. The L and P22 Erf proteins are very similar (89%) in their C-terminal 55 amino acids, and although their genome positions and very similar C-termini suggest similar functions, the two N-terminal regions are essentially unrelated (13% identical). Poteete *et al*. (Poteete, Sauer et al. 1983) showed that Erf is a two-domain protein and it appears that this sequence similarity pattern is explained by horizontal exchange of one of the domains. Similarly, *cIII* was shown to be interchangeable between L and P22 in spite of only 55.6% identity at protein level (Bezdek, Soska et al. 1970).

The P22 gene *24* protein antiterminates early left and early right transcription to allow full expression of those operons. The L gene *24* protein is 96% identical to that of P22 over the N-terminal 75 amino acids, but their C-terminal 25-35 amino acids (they are different lengths) are essentially unrelated. The fact that these two proteins are functionally interchangeable (Schicklmaier and Schmieger 1997) and that their *nutL* and *nutR* boxB target sites are apparently identical (not shown) support the notion that L and P22 gp24 proteins have similar target specificities and that, like phage λ N protein, their specificities are controlled by their N-terminal 75 amino acids (Cocozaki, Ghattas et al. 2008, Krupp, Said et al. 2019).

#### Primary prophage immunity

The L *immC* region exhibits conservation of gene position for P22 *c2*, *cro* and *c1*, but divergence in sequence compared to P22 (Figures 2 and 3C). This agrees with the known distinct repressor target specificities of P22 and L (Hilliker 1981). DNA binding specificity differences no doubt have their origin in the divergence of protein sequence: 50.4%, 19.4% and 48.4%, respectively for CII, Cro and CI proteins. The divergence is consistent with earlier DNA sequence analysis (Schicklmaier and Schmieger 1997); note that here our sequences correct a number of errors in that sequence.

#### Replication region

Similarly, the L replication module (red in Figures 2 and 3C) displays low similarity for P22 replication genes *18* and *12*. The L gene *18* protein may be a distant homolog of phage λ DNA replication initiation protein gpO, but it is only very distantly related in sequence (15% identical); it is 25% identical to P22 gp18. This group of proteins vary in sequence according to their origin binding specificities. The observation that the individual P22 replication genes have different specificity from those of L and cannot be substituted by the parallel individual L genes, but that the whole replication region can be substituted (Hilliker 1981) suggests that the L replication origin also lies within this region. The λ *O* protein binds to the replication origin, which consist of four ~20 bp inexact repeats (called “iterons”) lying near the center of the gene *O* coding region (Tsurimoto and Matsubara 1981). The P22 *18* gene contains four ~20 bp repeats that are different from those of λ, and the L gene *18* contains six imprecise repeats of a distinct eleven bp sequence (TGTCCAACGGA); this repeat region is likely the origin of L replication. Thus, these three distantly related *ori* binding proteins appear each to bind different iterons within their coding regions. The other L replication protein, the product of gene *12*, is 29% identical to that of P22. Thus, by homology its gp12 should also be a DnaB type helicase like that of P22 (Wickner 1984).

#### The *nin* region and late operon activator

The 3’-terminal part of the early right operon in λ includes the *ren, nin* and *Q* genes (P22 *nin* and *23* genes). The *nin* region of phage λ (genes *ren* through *Q*) contains 11 non-essential genes named *ren*, *ninA* through *ninH, orf221* and gene *23* (Cheng, Court et al. 1995), and in P22, the corresponding region comprises ten genes, *ninA, B, D, E, F, G, H, X, Y* and *Z* (Vander Byl and Kropinski 2000, Pedulla, Ford et al. 2003). In the L genome this region is composed of 14 genes (light green Figure 2). The functions of these L genes are largely unknown, but *ninB* (also known as the *orf* gene in phage λ) and *rusA* have putative functions in homologous recombination (Tarkowski, Mooney et al. 2002) and as a Holliday junction resolvase (Sharples, Chan et al. 1994, Mahdi, Sharples et al. 1996), respectively. The L *ninA, D, E, F, H* and *Z* genes are almost identical to P22 genes at the nucleotide level (Figure 3D), whereas *ninB* and *ninX* are more divergent, sharing only 48.4 and 70.9 % DNA identity, respectively. In all lambdoid phages the *nin* region genes are exceptionally tightly packed together, with the presence of numerous ATGA sequence motifs between genes which comprise overlapping translation terminator TGA codons for the upstream gene and initiator ATG codons for the downstream genes (Kröger and Hobom 1982, Cheng, Court et al. 1995).

In the lambdoid phages, the last gene in the early right operon encodes a protein that activates transcription of the late operon by antitermination of RNA polymerase (gene *Q* protein in λ). At least five quite different sequence types of these anti-terminators are known in lambdoid phages; these non-homologous yet functionally analogous proteins bind different target sequences (Grose and Casjens 2014, Yin, Kaelber et al. 2019). The λ and P22 proteins are very similar and can substitute for one another (type 1), but the L protein is very different from them (type 3; only 12% identical to that of P22) and 80% identical to its phage 21 homolog whose atomic structure and binding site are known (Yin, Kaelber et al. 2019). In spite of the gp23 similarity with 21 gpQ, the 21 Q binding element (QBE) sequence and the late operon start site are substantially different in L. In 21, QBE is two near-perfect tandem 8 bp copies of ATTGAGCA/AaTGAGCA (lower case marks non-identities) overlapping the 3’-end of gene *23*; in L there are two tandem very similar sequences of AATTATCC/AtTTAgCC (bp 37,761-37,776). It seems quite likely that this is the L QBE, and similarly the L late operon initiation site should be very near bp 37,791. Most of the differences between the 21 and L gpQ-like proteins are between amino acid 94 and the C-terminus; this divergent region is the segment of the 21 protein that contacts the QBE. Three of the six amino acids reported to contact the QBE in 21 (Yin, Kaelber et al. 2019) are different in the two phages, so different target specificities are not surprising.

#### The L late operon - tRNA genes

P22 does not carry any tRNA genes, but two tRNA genes lie near the 5’end of the L late operon (the first starts 46 bp downstream of putative late mRNA start discussed above). They appear to have anti-codons GUU and UAA, which should insert asparagine and leucine, respectively. Some, but not all P22-like phages are known to carry tRNA genes in this location; for example, *Shigella* phage Sf6 has asparagine and threonine tRNAs here (64). It is not known if these tRNA molecules might be excised from the late mRNA or expressed independently from a currently unidentified promoter(s). However, 24 of 25 codons specifying Asn in the major capsid gene (and hence the most abundantly expressed protein) are AAC (matching the phage tRNA’s GUU anticodon) suggesting that at least one of these tRNAs might be encoded to supplement the tRNA pool during intracellular development of the virion.

#### The L late operon - lysis genes

The L lysis module contains genes *13, 19, 15* and *Rz1* and lies in the late operon immediately downstream of the tRNA genes; as in all lambdoid phages where it has been studied, these genes are expressed as part of the late operon. Two of these four genes, overlapping *15* and *Rz1*, are subunits of a spanin that functions to disrupt the outer membrane (Kongari, Rajaure et al. 2018). These two proteins are quite similar (64% and 93% identical, respectively) to their P22 homologs. On the other hand, lysis genes *13* and *19*, which encode a putative holin and endolysin, respectively, are very different from their P22 counterparts (Figure 3D).

### Genome modularity and transcription

The clustering of genes with related functions in the P22-like phage genomes has been discussed (Vander Byl and Kropinski 2000, Mmolawa, Schmieger et al. 2003, Casjens, Winn-Stapley et al. 2004, Villafane, Zayas et al. 2008, Casjens and Thuman-Commike 2011, Hendrix and Casjens 2013), and phage L clearly conforms to this pattern. Indeed, comparison of L with P22 shows a very highly mosaic relationship in which at least 25 sections of very similar DNA are separated by unrelated patches (Figure 3). Apparent transcriptional units (as indicated by the positions of predicted terminators) do not coincide perfectly with known operons that have been defined in P22 (Figure 2). For example, a putative terminator is present at the right end of the procapsid-encoding gene module. This terminator lies within the putative late operon, which has only one promoter in P22, and it is almost certainly the case that the closely related L late operon also has only one promoter. Such intra-operon terminators are not unique to L, and their roles are not known. Possible consequences could include partial termination to reduce the amount of mRNA made for the downstream portion of the messenger RNA or they could stop spurious transcription events that initiate without gp23 or gp24 mediated antitermination.

### Phage L Virion proteins

LC MS/MS mass spectrometry analysis (as described in Material and Methods) of the proteins present in preparations CsCl density gradient purified L *cI*^−^40 *13*^−^*am*43 virions found evidence for the presence of 22 phage L-encoded proteins (Supplementary data protein excel file). Peptide fragments were observed that match all the virion assembly proteins whose products are known P22 virion components (*1, 5, 4, 10, 26, 14, 7, 20, 16* and *9* – listed in genetic map order). The L virion preparations also contain Dec protein, scaffolding protein and the small terminase subunit. Dec is specific to phage L and its close relatives (Gilcrease, Winn-Stapley et al. 2005, Casjens and Thuman-Commike 2011), and 180 molecules have been shown to be bound on the outside surface of the capsid (Tang, Gilcrease et al. 2006). Scaffolding protein gp8, the product of gene *8*, is also present in the L virion preparations, but its homolog is not known to be present in P22 virions. Scaffolding protein is present in large numbers in P22 procapsids but is completely absent from virions (Casjens and King 1974, King and Casjens 1974). It is unclear whether, unlike the situation in P22, some gp8 molecules might remain in L virions or whether this indicates a low level of contamination in the L virion preparations by virion precursor particles. Since the phage lysate preparation also contains host-derived cell envlope peptides (see Table S2), it’s likely that virion precursor particles are also present in the step-gradient cesium preparation. The 9 remaining proteins might fall in that category, especially DNA binding proteins (Cro, Int and gp18).

### DNA methylation and restriction

When phages are propagated in their bacterial host, they often acquire host-specified DNA modifications that protect them against cognate host-specified restriction endonuclease systems, or that mediate other epigenetic functions (Sánchez-Romero and Casadesus 2020). At least six methylation motifs are known in the propagation host, *Salmonella enterica* serovar Typhimurium LT2 ((Roberts, Vincze et al. 2015); REBASE Organism number 18099). Three motifs result from enterobacterial “orphan” methyltransferases (M; M.SenLT2 Dam, M.SenLT2 Dcm and M.SenLT2IV; (Roberts, Vincze et al. 2015)), and three are associated with restriction (RM) phenomena (M.SenLT2I, M.SenLT2II and M.SenLT2III). Two of these RM systems have been well characterized. M.SenLT2I (known in the literature as the StyLTI RM system (Colson and Colson 1971)) is a Type III enzyme, and M.SenLT2II (StySB or StyLTII (Fuller-Pace, Bullas et al. 1984)) is Type I. The third system, RM.SenLT2III (StySA), is known to confer m6A modification but its protection/restriction mechanism of action remains unclear (Hattman, Schlagman et al. 1976).

#### Number of RM sites

Historically, three *Salmonella* phages were used to score the activity of the three RM activities: P22, L and P3 (Bullas and Ryu 1983). This practice was due to the uneven magnitude of restriction of each phage by individual R activities: SenLT2I (StyLT) was highly efficient on all phages, with restriction of 10^3^-10^4^-fold; SenLT2II (StySB) did not restrict P22 at all, so P3 was used (or even λ, in intergeneric hybrids (Colson and Van Pel 1974)); and SenLT2III (StySA) restricted P22 about two-fold, but restricted L 100-fold (Colson and Colson 1971). While numerous anti-restriction activities are counteracted by phage and plasmid encoded machinery (Tock and Dryden 2005, Labrie, Samson et al. 2010), a simple evolutionary evasive strategy is the purging of sequences specifically recognized by endonucleases (Murray and Murray 1974, Rusinov, Ershova et al. 2018). In this instance we find that the number of sites for the various enzymes is consistent with this strategy. As shown in Table 2, both P22 and L have more than 40 sites at which SenLT2I (StyLT) should act. Both P22 and L have only one site for SenLT2II (StySB); the Type I enzyme class is known to require multiple sites for action (Loenen, Dryden et al. 2014). In addition, if the phage L *ral* function acts as does the λ homolog (by promoting Type I methylation of progeny genomes (Loenen, Dryden et al. 2014)), restriction would be further reduced. SenLT2III (StySA) is molecularly uncharacterized, but we observe that P22 has only three sites while L has 13.

**Table 2:**
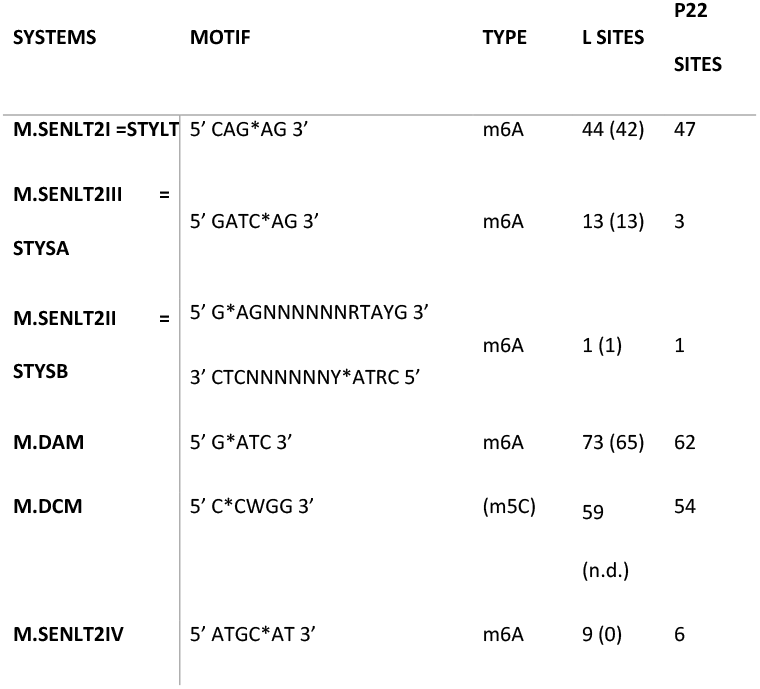
*S. enterica* serovar Typhimurium methyltransferase recognition sites present in the L genome. Column 1: the MTase name used in REBASE with the name most encountered in the literature. Column 2: recognition sequence, in which symmetric sites are shown once; asymmetric sites methylated only on one strand are shown once; asymmetric sites methylated on both strands are shown twice. Column 3: methylation type. Column 4 and 5: the number of methylation motif sequences present in L and P22 genomes. In parenthesis for L is the number of methylated sites observed by the PacBio sequence analysis; N.D. (not detected), m5C was not determined by this approach.

#### Undermethylation

In Table 2, we also find no modification of sites in the L genome for the normally-silent orphan methyltransferase M.SenLT2IV, as previously observed in *E. coli* and *S. enterica* chromosomal DNA (Broadbent, Balbontin et al. 2007). Partial modification at most Dam sites is consistent with findings of others; virion genomes of P22 and λ are undermethylated by Dam and Dcm, possibly because replication outstrips modification activity (Casjens, Hayden et al. 1983, Szyf, Avraham-Haetzni et al. 1984). Methylation of m5C by M.Dcm was not determined here, due to technical limitations of Pacific Bioscience long-read sequencing in detection of m5C (Flusberg, Webster et al. 2010).

Interestingly, we also observe selective undermethylation of specific sites; Dam methylation is completely absent at the four sites upstream of *gtrABC*, and also at one site within the body of the gene. The P22 version of this locus is known to be the target of Dam/OxyR-mediated phase variation in the prophage state (Broadbent, Davies et al. 2010). The host regulator OxyR blocks methylation at one pair of sites in one phase or at the other pair of sites in the other phase, resulting in variable transcription activity. Possibly OxyR plays a role during DNA packaging to prevent Dam modification in this region.

### Summary

Overall, these sequencing data confirm the P22-like L genomic organization previously observed through other less precise methods. It elucidated the highly mosaic relationship between L and P22 and discovered the extreme sequence similarity between phages L and ST64T even though the latter two phages were isolated 35 years apart. Sequence analysis showed that L is potentially more vulnerable to M.SenLT2III (StySA) restriction activity than P22 due to its greater number of recognition sites, making it a better tester for this molecularly uncharacterized genome-defense system. M.SenLT2IV methylation is absent, and Dam methylation incomplete, as also seen in λ and P22.

## ACKNOWLEDGEMENTS

We thank Eddie Gilcrease, Werner Bode and David Botstein for providing the L phage isolates and Anca Segall for kindly sharing the LB5000 *Salmonella* strain.

## ADDITIONAL MARTERIALS

Supplemental material available on GSA Figshare portal.

---

**Table S1: Annotations**

This excel file contains four sheets. The first one is a guide to read the file, second one is gathering the predictions from PHAST and RAST, third one, data from literature and the last one contains the predictions of rho-independent terminators.

**Table S2: Proteomic data**

This excel file contains the output of the proteomic analysis. The first sheet gathers the data from the spectral counts and the second one an analysis by peak era.

## REFERENCES ENDNOTE G3

Adhikari, P., G. Allison, B. Whittle and N. K. Verma (1999). “Serotype 1a O-antigen modification: molecular characterization of the genes involved and their novel organization in the Shigella flexneri chromosome.” J Bacteriol 181(15): 4711.–4718.

Arndt, D., J. R. Grant, A. Marcu, T. Sajed, A. Pon, Y. Liang and D. S. Wishart (2016). “PHASTER: a better, faster version of the PHAST phage search tool.” Nucleic Acids Research 44(W1): W16–W21.

Aziz, R. K., D. Bartels, A. Best, M. DeJongh, T. Disz, R. A. Edwards, K. Formsma, S. Gerdes, E. M. Glass, M. Kubal, F. Meyer, G. J. Olsen, R. Olson, A. L. Osterman, R. A. Overbeek, L. K. McNeil, D. Paarmann, T. Paczian, B. Parrello, G. D. Pusch, C. Reich, R. Stevens, O. Vassieva, V. Vonstein, A. Wilke and O. Zagnitko (2008). “The RAST Server: Rapid annotations using subsystems technology.” BMC Genomics 9.

Bezdek, M. and P. Amati (1967). “Properties of P22 and A related Salmonella typhimurium phage. I. General features and host specificity.” Virology 31(2): 272–278.

Bezdek, M. and P. Amati (1968). “Evidence for two immunity regulator systems in temperature bacteriophages P22 and L.” Virology 36(4): 701–703.

Bezdek, M., J. Soska and P. Amati (1970). “Properties of P22 and a related Salmonella typhimurium phage. III. Studies on clear-plaque mutants of phage L.” Virology 40(3): 505–513.

Bode, W. (1979). “Regulation of late functions in Salmonella bacteriophages P22 and L studied by assaying endolysin synthesis.” Journal of virology 32(1): 1–7.

Broadbent, S. E., R. Balbontin, J. Casadesus, M. G. Marinus and M. van der Woude (2007). “YhdJ, a nonessential CcrM-like DNA methyltransferase of Escherichia coli and Salmonella enterica.” J Bacteriol 189(11): 4325–4327.

Broadbent, S. E., M. R. Davies and M. W. van der Woude (2010). “Phase variation controls expression of Salmonella lipopolysaccharide modification genes by a DNA methylation-dependent mechanism.” Molecular Microbiology 77(2): 337–353.

Bullas, L. R. and J. I. Ryu (1983). “Salmonella typhimurium LT2 strains which are r-m+ for all three chromosomally located systems of DNA restriction and modification.” J Bacteriol 156(1): 471–474.

Casjens, S., M. Hayden, E. Jackson and R. Deans (1983). “Additional restriction endonuclease cleavage sites on the bacteriophage P22 genome.” Journal of virology 45(2): 864–867.

Casjens, S. and J. King (1974). “P22 morphogenesis I: Catalytic scaffolding protein in capsid assembly.” Journal of Supramolecular Structure 2(2-4): 202–224.

Casjens, S., D. A. Winn-Stapley, E. B. Gilcrease, R. Morona, C. Kühlewein, J. E. H. Chua, P. A. Manning, W. Inwood and A. J. Clark (2004). “The chromosome of Shigella flexneri bacteriophage Sf6: complete nucleotide sequence, genetic mosaicism, and DNA packaging.” Journal of Molecular Biology 339(2): 379–394.

Casjens, S. R. and E. B. Gilcrease (2009). “Determining DNA packaging strategy by analysis of the termini of the chromosomes in tailed-bacteriophage virions.” Methods in molecular biology (Clifton, NJ) 502: 91–111.

Casjens, S. R. and P. A. Thuman-Commike (2011). “Evolution of mosaically related tailed bacteriophage genomes seen through the lens of phage P22 virion assembly.” Virology 411(2): 393–415.

Chan, P. P. and T. M. Lowe (2019). “tRNAscan-SE: Searching for tRNA Genes in Genomic Sequences.” Methods Mol Biol 1962: 1–14.

Cheng, S. W., D. L. Court and D. I. Friedman (1995). “Transcription termination signals in the nin region of bacteriophage lambda: identification of Rho-dependent termination regions.” Genetics 140(3): 875–887.

Clark, T. A., I. A. Murray, R. D. Morgan, A. O. Kislyuk, K. E. Spittle, M. Boitano, A. Fomenkov, R. J. Roberts and J. Korlach (2012). “Characterization of DNA methyltransferase specificities using single-molecule, real-time DNA sequencing.” Nucleic Acids Research 40(4): e29–e29.

Cocozaki, A. I., I. R. Ghattas and C. A. Smith (2008). “Bacteriophage P22 antitermination boxB sequence requirements are complex and overlap with those of lambda.” Journal of Bacteriology 190(12): 4263–4271.

Colson, C. and A. M. Colson (1971). “A new Salmonella typhimurium DNA host specificity.” J Gen Microbiol 69(3): 345–351.

Colson, C. C. and A. A. Van Pel (1974). “DNA restriction and modification systems in Salmonella. I. SA and SB, two Salmonella typhimurium systems determined by genes with a chromosomal location comparable to that of the Escherichia coli hsd genes.” Molecular Genetics and Genomics 129(4): 325–337.

Earnshaw, W., S. Casjens and S. C. Harrison (1976). “Assembly of the head of bacteriophage P22: x-ray diffraction from heads, proheads and related structures.” Journal of Molecular Biology 104(2): 387–410.

Favre, R., P. Amati and M. Bezdek (1968). “Properties of P22 and a related Salmonella typhimurium phage. II. Effect of H markers on infection of strain 1559.” Virology 35(2): 238–247.

Flusberg, B. A., D. R. Webster, J. H. Lee, K. J. Travers, E. C. Olivares, T. A. Clark, J. Korlach and S. W. Turner (2010). “Direct detection of DNA methylation during single-molecule, real-time sequencing.” Nat Methods 7(6): 461–465.

Fuller-Pace, F. V., L. R. Bullas, H. Delius and N. E. Murray (1984). “Genetic recombination can generate altered restriction specificity.” Proc. Natl. Acad. Sci. USA 81: 6095–6099.

Gilcrease, E. B., D. A. Winn-Stapley, F. C. Hewitt, L. Joss and S. R. Casjens (2005). “Nucleotide sequence of the head assembly gene cluster of bacteriophage L and decoration protein characterization.” J Bacteriol 187(6): 2050–2057.

Gordon, D., C. Abajian and P. Green (1998). “Consed: a graphical tool for sequence finishing.” Genome Research 8(3): 195–202.

Grose, J. H. and S. R. Casjens (2014). “Understanding the enormous diversity of bacteriophages: the tailed phages that infect the bacterial family Enterobacteriaceae.” Virology 468-470: 421–443.

Hattman, S., S. Schlagman, L. Goldstein and M. Frohlich (1976). “Salmonella typhimurium SA host specificity system is based on deoxyribonucleic acid-adenine methylation.” J Bacteriol 127(1): 211–217.

Hayden, M., M. B. Adams and S. Casjens (1985). “Bacteriophage L: Chromosome physical map and structural proteins.” Virology 147(2): 431–440.

Hendrix, R. W. and S. Casjens (2013). Bacteriophage lambda and its Genetic Neighborhood. R. Calendar, Oxford University Press: 409–477.

Hilliker, S. (1981). “Characterization of the Salmonella Phage L early genes using lambda imm L hybrid phages.” Virology 114(1): 161–174.

Hunt, M., N. D. Silva, T. D. Otto, J. Parkhill, J. A. Keane and S. R. Harris (2015). “Circlator: automated circularization of genome assemblies using long sequencing reads.” Genome Biol 16: 294.

Kahmann, R. and H. Prell (1971). “Complementation between P22 amber mutants and phage L.” Mol Gen Genet 113(4): 363–366.

King, J. and S. Casjens (1974). “Catalytic head assembling protein in virus morphogenesis.” Nature 251(5471): 112–119.

Kongari, R., M. Rajaure, J. Cahill, E. Rasche, E. Mijalis, J. Berry and R. Young (2018). “Phage spanins: diversity, topological dynamics and gene convergence.” BMC Bioinformatics 19(1): 326–326.

Koren, S., B. P. Walenz, K. Berlin, J. R. Miller, N. H. Bergman and A. M. Phillippy (2017). “Canu: scalable and accurate long-read assembly via adaptive k-mer weighting and repeat separation.” Genome Res 27(5): 722–736.

Krey, J. F., P. A. Wilmarth, J.-B. Shin, J. Klimek, N. E. Sherman, E. D. Jeffery, D. Choi, L. L. David and P. G. Barr-Gillespie (2014). “Accurate label-free protein quantitation with high- and low-resolution mass spectrometers.” Journal of Proteome Research 13(2): 1034–1044.

Kröger, M. and G. Hobom (1982). “A chain of interlinked genes in the ninR region of bacteriophage lambda.” Gene 20(1): 25–38.

Krupp, F., N. Said, Y.-H. Huang, B. Loll, J. Bürger, T. Mielke, C. M. T. Spahn and M. C. Wahl (2019). “Structural Basis for the Action of an All-Purpose Transcription Anti-termination Factor.” Molecular cell 74(1): 143–157.e145.

Labrie, S. J., J. E. Samson and S. Moineau (2010). “Bacteriophage resistance mechanisms.” Nat Rev Microbiol 8(5): 317–327.

Langmead, B. and S. L. Salzberg (2012). “Fast gapped-read alignment with Bowtie 2.” Nat Methods 9(4): 357–359.

Leavitt, J. C., E. B. Gilcrease, K. Wilson and S. R. Casjens (2013). “Function and horizontal transfer of the small terminase subunit of the tailed bacteriophage Sf6 DNA packaging nanomotor.” Virology 440(2): 117–133.

Li, H., B. Handsaker, A. Wysoker, T. Fennell, J. Ruan, N. Homer, G. Marth, G. Abecasis, R. Durbin and S. Genome Project Data Processing (2009). “The Sequence Alignment/Map format and SAMtools.” Bioinformatics 25(16): 2078–2079.

Lindsey, D. F., C. Martínez and J. R. Walker (1992). “Physical map location of the Escherichia coli attachment site for the P22 prophage (attP22).” Journal of Bacteriology 174(11): 3834–3835.

Loenen, W. A., D. T. Dryden, E. A. Raleigh and G. G. Wilson (2014). “Type I restriction enzymes and their relatives.” Nucleic Acids Res 42(1): 20–44.

Lowe, T. M. and P. P. Chan (2016). “tRNAscan-SE On-line: integrating search and context for analysis of transfer RNA genes.” Nucleic Acids Res 44(W1): W54–57.

Mahdi, A. A., G. J. Sharples, T. N. Mandal and R. G. Lloyd (1996). “Holliday junction resolvases encoded by homologous rusA genes in Escherichia coli K-12 and phage 82.” Journal of Molecular Biology 257(3): 561–573.

McClelland, M., K. E. Sanderson, J. Spieth, S. W. Clifton, P. Latreille, L. Courtney, S. Porwollik, J. Ali, M. Dante, F. Du, S. Hou, D. Layman, S. Leonard, C. Nguyen, K. Scott, A. Holmes, N. Grewal, E. Mulvaney, E. Ryan, H. Sun, L. Florea, W. Miller, T. Stoneking, M. Nhan, R. Waterston and R. K. Wilson (2001). “Complete genome sequence of Salmonella enterica serovar Typhimurium LT2.” Nature 413(6858): 852–856.

Mmolawa, P. T., H. Schmieger, C. P. Tucker and M. W. Heuzenroeder (2003). “Genomic structure of the Salmonella enterica serovar Typhimurium DT 64 bacteriophage ST64T: Evidence for modular genetic architecture.” Journal of Bacteriology 185(11): 3473–3475.

Mosley, A. L., M. E. Sardiu, S. G. Pattenden, J. L. Workman, L. Florens and M. P. Washburn (2011). “Highly reproducible label free quantitative proteomic analysis of RNA polymerase complexes.” Molecular & cellular proteomics: MCP 10(2): M110.000687.

Murray, N. E. and K. Murray (1974). “Manipulation of restriction targets in phage lambda to form receptor chromosomes for DNA fragments.” Nature 251(5475): 476–481.

Pedulla, M. L., M. E. Ford, J. M. Houtz, T. Karthikeyan, C. Wadsworth, J. A. Lewis, D. Jacobs-Sera, J. Falbo, J. Gross, N. R. Pannunzio, W. Brucker, V. Kumar, J. Kandasamy, L. Keenan, S. Bardarov, J. Kriakov, J. G. Lawrence, W. R. Jacobs, Jr., R. W. Hendrix and G. F. Hatfull (2003). “Origins of highly mosaic mycobacteriophage genomes.” Cell 113(2): 171–182.

Pedulla, M. L., M. E. Ford, T. Karthikeyan, J. M. Houtz, R. W. Hendrix, G. F. Hatfull, A. R. Poteete, E. B. Gilcrease, D. A. Winn-Stapley and S. R. Casjens (2003). “Corrected sequence of the bacteriophage p22 genome.” J Bacteriol 185(4): 1475–1477.

Poteete, A. R., R. T. Sauer and R. W. Hendrix (1983). “Domain structure and quaternary organization of the bacteriophage P22 Erf protein.” Journal of Molecular Biology 171(4): 401418.

Roberts, R. J., T. Vincze, J. Posfai and D. Macelis (2015). “REBASE--a database for DNA restriction and modification: enzymes, genes and genomes.” Nucleic Acids Res 43(Database issue): D298299.

Rusinov, I. S., A. S. Ershova, A. S. Karyagina, S. A. Spirin and A. V. Alexeevski (2018). “Avoidance of recognition sites of restriction-modification systems is a widespread but not universal anti-restriction strategy of prokaryotic viruses.” BMC Genomics 19(1): 885.

Sánchez-Romero, M. A. and J. Casadesus (2020). “The bacterial epigenome.” Nature Reviews Microbiology 18(1): 7–20.

Sauer, R. T., W. Krovatin, J. DeAnda, P. Youderian and M. M. Susskind (1983). “Primary structure of the immI immunity region of bacteriophage P22.” Journal of Molecular Biology 168(4): 699713.

Schicklmaier, P. and H. Schmieger (1997). “Sequence comparison of the genes for immunity, DNA replication, and cell lysis of the P22-related Salmonella phages ES18 and L.” Gene 195(1): 93–100.

Sharples, G. J., S. N. Chan, A. A. Mahdi, M. C. Whitby and R. G. Lloyd (1994). “Processing of intermediates in recombination and DNA repair: identification of a new endonuclease that specifically cleaves Holliday junctions.” The EMBO journal 13(24): 6133–6142.

Susskind, M. M. and D. Botstein (1978). “Repression and immunity in Salmonella phages P22 and L: phage L lacks a functional secondary immunity system.” Virology 89(2): 618–622.

Susskind, M. M., D. Botstein and A. Wright (1974). “Superinfection exclusion by P22 prophage in lysogens of Salmonella typhimurium. III. Failure of superinfecting phage DNA to enter sieA+ lysogens.” Virology 62(2): 350–366.

Susskind, M. M., A. Wright and D. Botstein (1974). “Superinfection exclusion by P22 prophage in lysogens of Salmonella typhimurium. IV. Genetics and physiology of sieB exclusion.” Virology 62(2): 367–384.

Szyf, M., K. Avraham-Haetzni, A. Reifman, J. Shlomai, F. Kaplan, A. Oppenheim and A. Razin (1984). “DNA methylation pattern is determined by the intracellular level of the methylase.” Proceedings of the National Academy of Sciences of the United States of America 81(11): 32783282.

Tang, L., E. B. Gilcrease, S. R. Casjens and J. E. Johnson (2006). “Highly discriminatory binding of capsid-cementing proteins in bacteriophage L.” Structure/Folding and Design 14(5): 837–845.

Tarkowski, T. A., D. Mooney, L. C. Thomason and F. W. Stahl (2002). “Gene products encoded in the ninR region of phage lambda participate in Red-mediated recombination.” Genes to cells: devoted to molecular & cellular mechanisms 7(4): 351–363.

Tock, M. R. and D. T. Dryden (2005). “The biology of restriction and anti-restriction.” Curr Opin Microbiol 8(4): 466–472.

Tsurimoto, T. and K. Matsubara (1981). “Purified bacteriophage lambda O protein binds to four repeating sequences at the lambda replication origin.” Nucleic Acids Research 9(8): 1789–1799.

Vander Byl, C. and A. M. Kropinski (2000). “Sequence of the genome of Salmonella bacteriophage P22.” J Bacteriol 182(22): 6472–6481.

Villafane, R., M. Zayas, E. B. Gilcrease, A. M. Kropinski and S. R. Casjens (2008). “Genomic analysis of bacteriophage epsilon 34 of Salmonella enterica serovar Anatum (15+).” BMC Microbiology 8: 227.

Wick, R. R., L. M. Judd, C. L. Gorrie and K. E. Holt (2017). “Unicycler: Resolving bacterial genome assemblies from short and long sequencing reads.” PLoS Comput Biol 13(6): e1005595.

Wickner, S. (1984). “Oligonucleotide synthesis by Escherichia coli dnaG primase in conjunction with phage P22 gene 12 protein.” Journal of Biological Chemistry 259(22): 14044–14047.

Wiggins, B. A. and S. Hilliker (1985). “Map of DNA homology between the genomes of Salmonella bacteriophages P22 and L.” Journal of virology 56(3): 1034–1036.

Winston, F., D. Botstein and J. H. Miller (1979). “Characterization of amber and ochre suppressors in Salmonella typhimurium.” Journal of Bacteriology 137(1): 433–439.

Wisniewski, J. R., A. Zougman, N. Nagaraj and M. Mann (2009). “Universal sample preparation method for proteome analysis.” Nat Methods 6(5): 359–362.

Wu, H., L. Sampson, R. Parr and S. Casjens (2002). “The DNA site utilized by bacteriophage P22 for initiation of DNA packaging.” Mol Microbiol 45(6): 1631–1646.

Yin, Z., J. T. Kaelber and R. H. Ebright (2019). “Structural basis of Q-dependent antitermination.” Proceedings of the National Academy of Sciences.

Zhou, Y., Y. Liang, K. H. Lynch, J. J. Dennis and D. S. Wishart (2011). “PHAST: A Fast Phage Search Tool.” Nucleic Acids Research 39(SUPPL. 2): W347–352.

Zinder, N. D. and J. Lederberg (1952). “Genetic exchange in Salmonella.” Journal of Bacteriology 64(5): 679–699.

